# Analysis of the Modulation of RAF Signaling by 14-3-3 Proteins

**DOI:** 10.1101/2024.07.16.603736

**Authors:** Peter Carlip, Edward C. Stites

**Affiliations:** Department of Laboratory Medicine, Yale Cancer Center, Yale School of Medicine, 300 George St., New Haven, 06512, CT, USA

**Keywords:** Cell Signaling, Signaling Mechanisms, RAF, kinases, Biochemical Networks

## Abstract

The regulation of cellular biochemical signaling reactions includes the modulation of protein activity through a variety of processes. For example, signaling by the RAF kinases, which are key transmitters of extracellular growth signals downstream from the RAS GTPases, is modulated by dimerization, protein conformational changes, post-translational modifications, and protein-protein interactions. 14-3-3 proteins are known to play an important role in RAF signal regulation, and have the ability to stabilize both inactive (monomeric) and active (dimeric) states of RAF. It is poorly understood how these antagonistic roles ultimately modulate RAF signaling. To investigate, we develop a mathematical model of RAF activation with both roles of 14-3-3, perform algebraic and numeric analyses, and compare with available experimental data. We derive the conditions necessary to explain experimental observations that 14-3-3 overexpression activates RAF, and we show that strong binding of 14-3-3 to Raf dimers alone is not generally sufficient to explain this observation. Our integrated analysis also suggests that RAF–14-3-3 binding is relatively weak for the reasonable range of parameter values, and suggests the Raf dimer–14-3-3 interactions are stabilized primarily by avidity. Lastly we find that in the limit of paired weak/avidity driven interactions between RAF and 14-3-3, the paired binding interactions may be reasonably approximated with a strong, single, equilibrium reaction. Overall, our work presents a mathematical model that can serve as a foundational piece for future, extended, studies of signaling reactions involving regulated RAF kinase activity.

## 1 Introduction

Many biological functions are controlled by the outputs of protein networks. For example, cellular proliferation can be regulated by the RAS-RAF-MEK-ERK network. There are three RAF proteins in humans: ARAF, BRAF, and CRAF, which are encoded by *ARAF, BRAF*, and *RAF1* genes respectively, as well as the closely related KSR1 and KSR2 proteins which are encoded by *KSR1* and *KSR2*. In response to various stimuli, the RAS proteins (KRAS, NRAS, and HRAS) are activated, and the activation of RAS leads to the activation of RAF which in turn activates MEK to propagate the signal onward. Although this basic concept of a signal propagating down a cascade of reactions is appealing and intuitive, actual signaling is more complex both with respect to the organization of the network and to the steps that influence whether a protein is in an “active” form that can further propagate signals or is in an “inactive” form and cannot. The activation of RAF proteins, in particular, is quite complicated (Lavoie and Therrien, 2015).

Although significant progress has been made in determining the steps involved in RAF activation, many aspects of the process remain poorly understood. For example, the modulation of RAF signals by 14-3-3 proteins is well-appreciated to occur but incompletely understood. (There are 7 different 14-3-3 proteins: 14-3-3 *β, ε, γ, η, σ, θ* (or *τ* in some sources), and ζ, coded by the genes *YWHAB, YWHAE, YWHAH, YWHAG, SFN, YWHAQ*, and *YWHAZ* respectively. These isoforms form homodimers and heterodimers in various combinations. For simplicity, we refer to any 14-3-3 dimer as 14-3-3.) It has been shown that 14-3-3 binds RAF at two separate phosphoserine sites: serine 729 (S729) and serine 365 (S365) in BRAF (Simanshu and Morrison, 2022), which correspond to serine 621 and serine 259 respectively in CRAF (Morrison et al, 1993). (The BRAF numbering is used throughout). 14-3-3 binding at S365 is inhibitory and binding at S729 is activating (Morrison et al, 1993; Tzivion et al, 1998). This is due to the two roles played by 14-3-3 binding: stabilization of closed, inactive RAF monomers,Park et al (2019) which involves binding at both S365 and S729, and stabilization of (potentially active) RAF dimers, involving binding solely at S729 of each monomer (Park et al, 2019; Martinez Fiesco et al, 2022). Fundamental aspects of these interactions remain incompletely defined; for example, measurements of the affinities of these bindings have varied widely (Muslin et al, 1996; Tzivion et al, 1998; Ghosh et al, 2015; Zhang et al, 2021), and the degree of avidity, or enhanced binding due to multivalent interactions (Erlendsson and Teilum, 2021), remains unclear.

In separate studies, we developed a mathematical model of RAF signaling that includes dimerization, conformational autoinhibition, and RAF inhibitor binding (Mendiratta and Stites, 2023), and then used that model to investigate the regulation of RAF signals by 14-3-3 (Mendiratta et al, 2021). That modeling suggested that the stabilization of the conformationally inactive form of RAF by 14-3-3 proteins can amplify the phenomena of “paradoxical activation,” in which RAF inhibitors actually cause increases in RAF signaling (Kholodenko, 2015; Rukhlenko et al, 2018; Fröhlich et al, 2023; Mendiratta and Stites, 2023). Our interpretation was also tested and supported by experimental results.

In order to create a tractable model of RAF regulation by 14-3-3 and the modulation of RAF inhibitor drug responses, the work in Mendiratta and Stites (2023) and Mendiratta et al (2021) necessarily simplified many aspects of RAF activation, including details of RAF–14-3-3 interactions. One such simplification was to only consider 14-3-3 binding to RAF monomers or dimers as a single equilibrium reaction between a 14-3-3 complex with both phosphorylated sites on RAF monomer (S365 and S729) or on a RAF dimer (S729 on each protomer in the dimer), rather than modeling the sites independently. It is not clear whether or how this simplification impacts model-based insights into 14-3-3 regulation of RAF.

With 14-3-3 a known, critical, modulator of RAF signals, we desired to study the effects of 14-3-3 on RAF activation without the previously used simplification. We constructed a model of RAF dimerization and the interaction of 14-3-3 with monomeric and dimeric forms of RAF. In order to make this model tractable, we did not include RAF inhibitor binding. Our new model can be solved analytically, allowing us to take derivatives of concentrations with respect to free 14-3-3 concentration. By evaluating values within a reasonable physiological range, we conclude that 14-3-3 binding to RAF is most likely avidity driven. By evaluating the Mendiratta et al (2021) transfection experiments, we also find that the observation that increased 14-3-3 increases dimerization, which combined with our new modeling suggests that only double-binding of 14-3-3 to RAF is significant. This work further substantiates earlier work, by justifying a critical simplification of Mendiratta et al (2021), and provides a foundational building-block that can be used to construct extended models of RAS/RAF and/or of growth factor signaling networks.

## 2 Model

### 2.1 Description

In this model of RAF–14-3-3 interactions, RAF can be in one of two conformations: open ([*O*]) and closed ([*C*]). When open, RAF may bind to 14-3-3 at the site S729, producing [729*O*]. It may also dimerize when open, whether bound to 14-3-3 at S729 or unbound (though only one protomer may be bound initially), producing [*dim*1] and [*dim*] respectively. RAF may also bind 14-3-3 when dimerized ([*dim*] → [*dim*1]). In addition, when a RAF dimer is bound to 14-3-3 at S729 of one of the component protomers, the second S729 may bind as well ([*dim*1] → [*dim*2]).

In the closed conformation, meanwhile, RAF can bind 14-3-3 at either S365 or S729, producing [365] and [729] respectively. When one site is bound, the other may also bind, producing [365/729]. RAF may close from either [*O*] or [729*O*], producing [*C*] and [729] respectively, but may not open when bound at S365. All binding to 14-3-3 in either state requires free 14-3-3 ([14-3-3]). These definitions are listed in A.1.

Within the model, all reactions are reversible, and detailed balance is maintained (see A.3), allowing for the system of eleven reactions to be described in eight equations, which are listed below with initial substitutions (equations 1-11). All equilibrium constants are in the form *k*_forward_/*k*_reverse_, with the forward direction described above (see A.2 for a list of definitions). For all binding interactions, this means they are in *K*_*A*_ form (*k*_*on*_/*k*_*off*_). The model is also shown graphically in Figure 1. The phosphorylation state of the serine residues is not modeled, and other aspects of RAF activation are abstracted into various rate constants (e.g. RAS binding as part of what determines the open:closed balance). This allows for a general, solvable model and accounts for uncertainty in the exact order of some aspects of RAF activation, but also means the model’s rate constants do not always correspond perfectly with the rates of specific chemical reactions. *K*_*ai*_, for instance, is not the ratio of the physical closing to opening rates. It expresses the ratio of concentrations in the open and closed states, which is affected by the rate of RAS binding and possibly inhibitory phosphorylation of RAF by ERK.

**Fig. 1:**
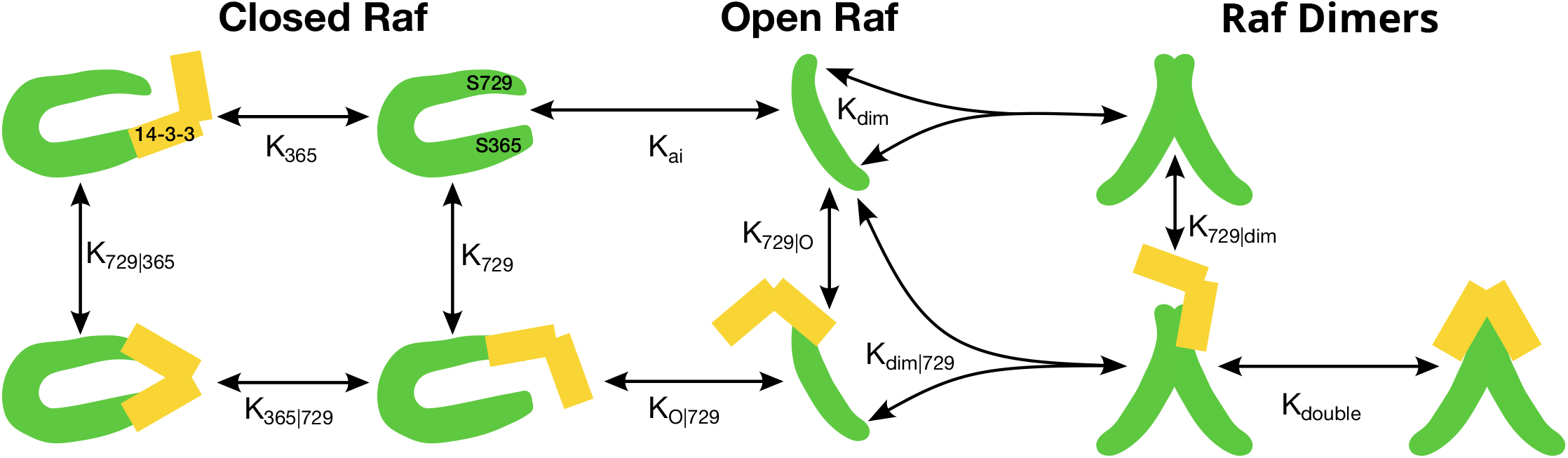
The RAF–14-3-3 model, including conformational changes, dimerization, and independent 14-3-3 binding at multiple RAF sites. The eleven reactions shown reduce to eight equilibrium equations, as each of the three cycles (S729 and S365 attachment to closed monomer, S729 attachment and opening, and S729 attachment and dimerization) allows for a simplification from detailed balance

### 2.2 Substitutions

To solve this system, we take an expression for the conserved total concentration of RAF as a sum of its states, and replace as many states as possible with expressions in terms of a single state ([*O*]) using the equilibrium relations in A.2:

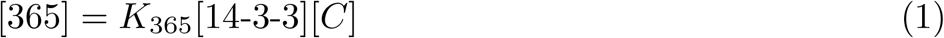

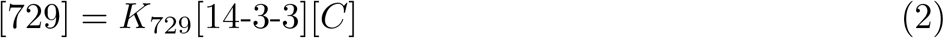

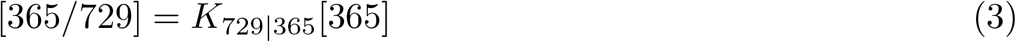

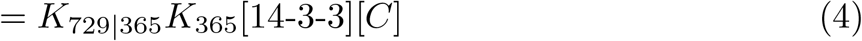

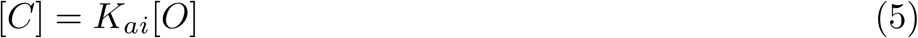

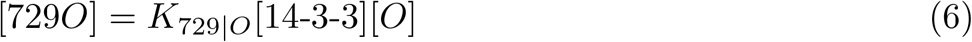

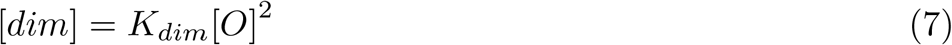

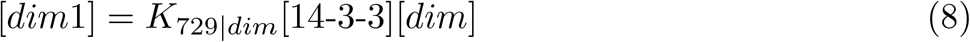

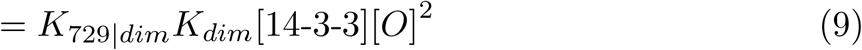

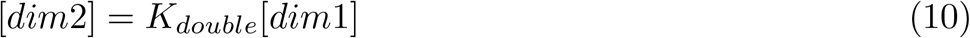

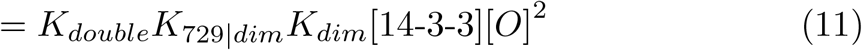

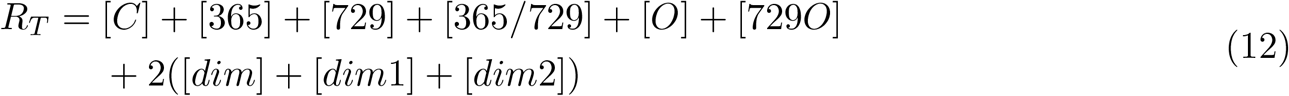

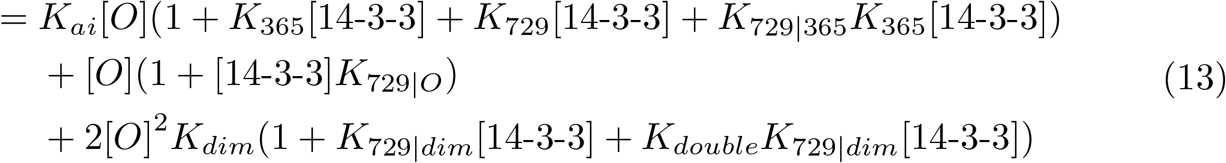

This system is quadratic in [*O*], given a constant free 14-3-3, and so it allows a solution for [*O*] using just the quadratic formula. As total 14-3-3 increases monotonically with free 14-3-3 (*∂*[Total 14-3-3]/*∂*[Free 14-3-3] > 0), this is just a rescaled version of the system with constant total RAF and 14-3-3. It is also possible to calculate total 14-3-3 once the system is solved with a given free 14-3-3, and use the result for numeric graphs with respect to total 14-3-3. In addition, dimers always become more common when more RAF is added or affinity of 14 –3-3 to dimers (*K* _729|*dim*_ or *K*_*double*_) is increased and become less common when RAF tends towards closed (increased *K*_*ai*_, e.g. because of reduced RAS activation).

From this solution, it is possible both to assess the system analytically and to numerically solve it for a given set of concentrations and equilibrium constants. Both are shown below.

## 3 Numerical Results

While the analytical solution derived in Section 2 allows us to determine the concentration of RAF states from the binding constants that serve as model parameters, many of those binding constants are effectively unknown, are incompletely defined, or vary between cell types. The total concentration of RAF can be estimated as ∼110 nM based on the results in Kulak et al (2014) and Fujioka et al (2006), and total con-centration of 14-3-3 can be estimated as well (see Sec. B). This analysis, which naively gives ∼100*×* as much 14-3-3 ζ and *η* alone (out of the 7 isoforms) as RAF, is com-plicated by the fact that 14-3-3 plays many other roles throughout the cell (Fu et al, 2000), making it unclear how much is available for interactions with RAF. The values of equilibrium constants are even less clear. For instance, reported binding affinities of 14-3-3 to target sites on RAF range from covalent-like energy differences in Zhang et al (2021) of hundreds of kcal/mol (from molecular dynamics simulations), to peptide affinities from Ghosh et al (2015) of ∼17 µM.

To analyze this model given these uncertainties, we conducted a variety of one– and two-variable parameter sweeps, taking as output the total concentration of RAF dimers (as only dimers are competent to signal (Hu et al, 2013)). A few patterns emerged throughout the parameter sweeps. Unsurprisingly, increasing *K*_365_ decreased total dimer concentration (see Figure 2a), as it causes 14-3-3 to more strongly stabilize closed monomers. Varying bonding constants to S729 (*K*_729_, *K*_729|*O*_, *K*_729|*dim*_) together (Figure 2b) has a more complex effect: increasing the affinity decreased unbound dimers and increased bound dimers, moving the system from one activation level to another. At the parameter values considered in Figure 2b and Figure 3b the bound dimer equilibrium was more active than the free dimer equilibrium, but it’s not clear that that would hold throughout all of parameter space. For instance, they must converge at sufficiently low 14-3-3 that few bound dimers can be formed regardless of affinity. In addition, increasing 14-3-3 concentration tends to reduce total dimer concentration, as can be seen by comparing the scales of Figure 2.

**Fig. 2:**
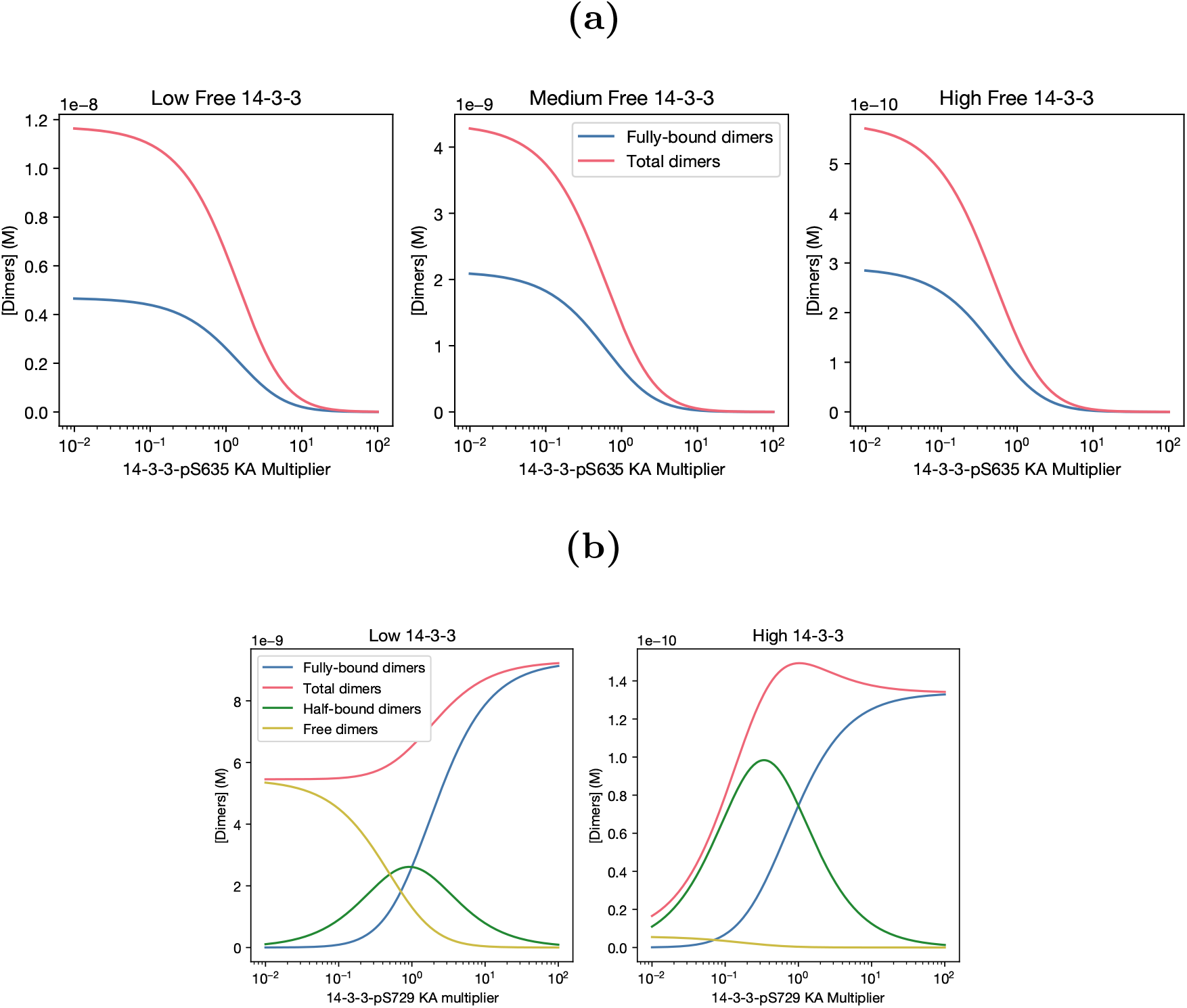
**(a)** At all 14-3-3 levels, increasing the strength of 14-3-3–S365 binding pulls more RAF to the closed conformation and reduces activation. **(b)** Strengthening 14-3-3–S729 binding moves dimers (and monomers) towards bound complexes, and tends to somewhat increase dimer formation. An equilibrium exists at both strong binding and fully bound RAF and at weak binding and fully unbound RAF

**Fig. 3:**
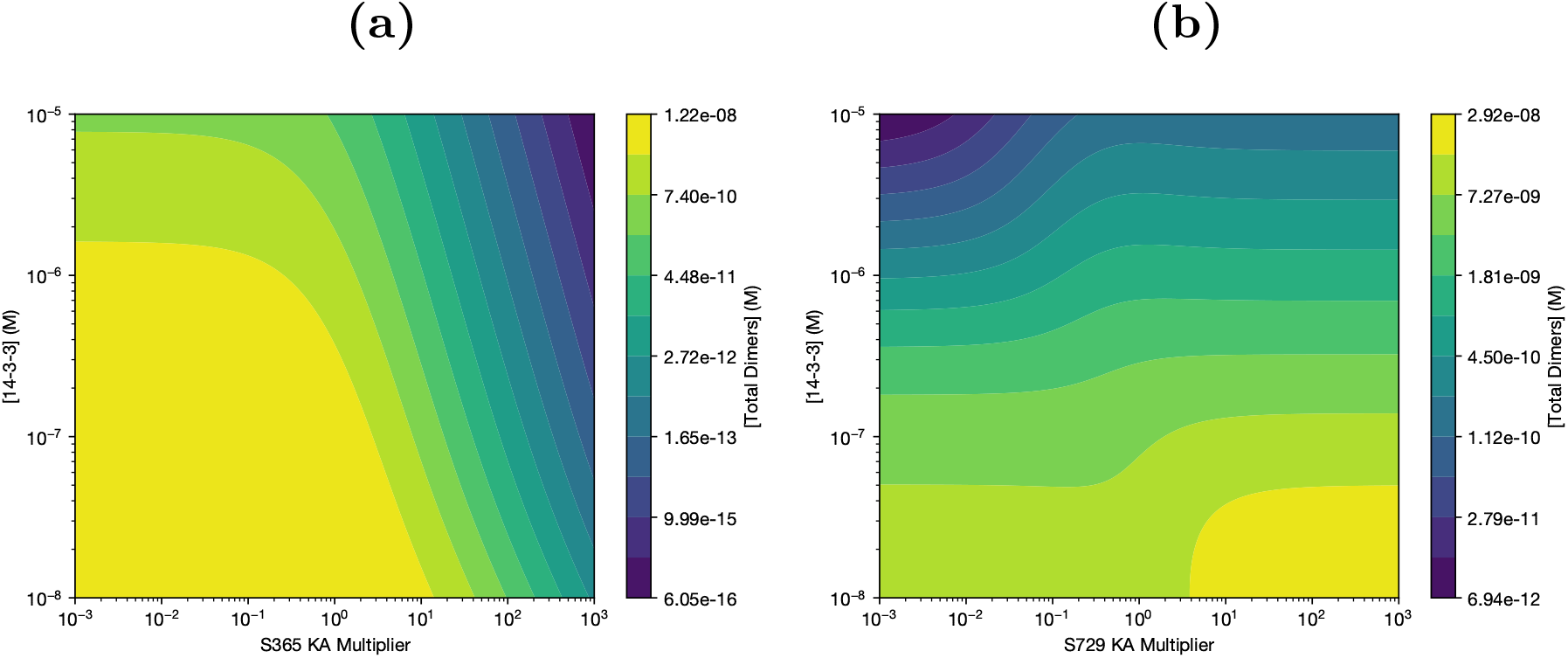
Contours of responses to 14-3-3 and Serine-specific affinity changes. **(a)** Increasing S365 affinity decreases activation at all 14-3-3 levels, not just the ones sampled in Figure 2a **(b)** The rise in activation corresponding to a change from unbound to bound dimers (see Figure 2b) is visible at various 14-3-3 concentrations. In either state, increasing 14-3-3 tends to decrease dimer levels

The parameter sweeps of Figure 2 are limited by the discrete set of 14-3-3 concentrations considered, so we also ran simultaneous dose-response plots of both 14-3-3 concentration and equilibrium constants, generating contour plots of dimer concentration (Figure 3). We see the patterns of response to equilibrium constants familiar from Figure 2: decreased dimerization in response to *K*_365_ (Figure 3a) and 2 equilibria at high and low S729 affinities (Figure 3b). It also becomes increasingly clear that, within the chosen parameter ranges (14-3-3 *K*_*D*_ around 100 nM, double-binding *K*_*D*_ around 1, 14-3-3 concentrations between 100 nM and 10 µM), increasing 14-3-3 will decrease dimer concentration.

This can then be extended to different combinations of affinities, as in Figure 4. Interestingly, as we see in Figure 4b, this pattern is not simply due to weak double binding. Increasing avidity to dimers (*K*_*double*_) increases dimer concentration, but it does not significantly affect the response of dimer concentration to 14-3-3. Instead, to expand the range of 14-3-3 concentration where increasing 14-3-3 increases dimers it is necessary to reduce the strength of single bonds, either alone (as in Figure 4c) or in combination with double binding (as in Figure 4a). This is not particularly intuitive, and analytic work with the solution was necessary to understand the mechanism leading to this behavior.

**Fig. 4:**
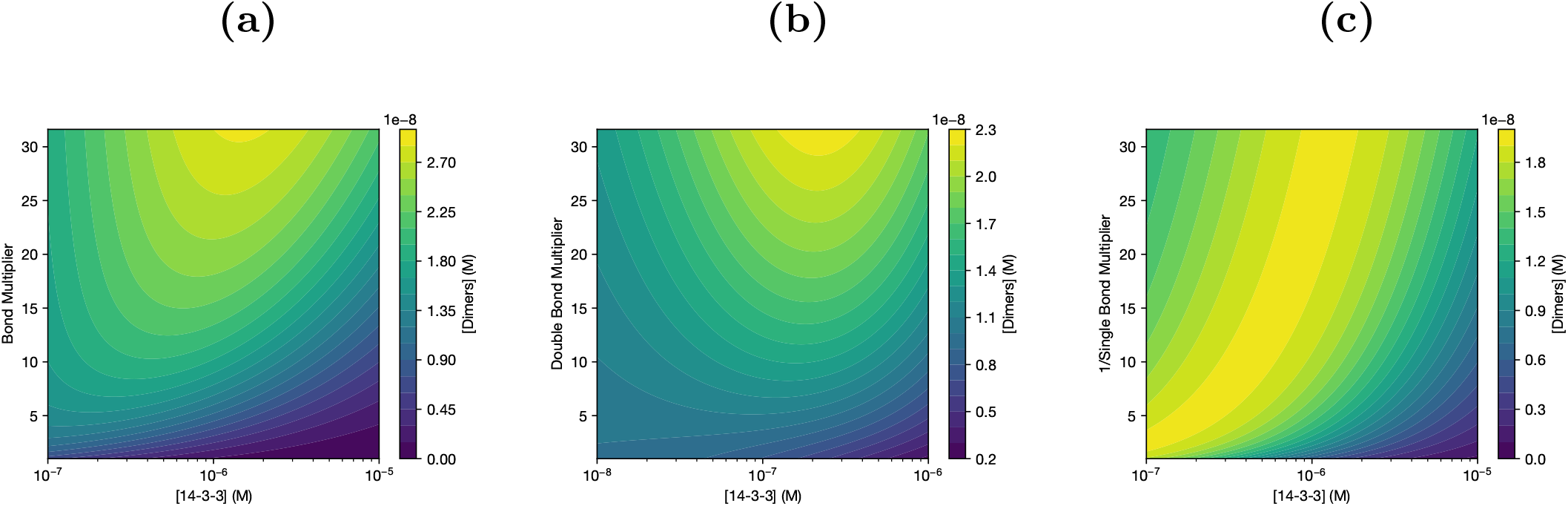
Contour plots of dimers vs [14-3-3] and equilibrium constants. **(a)** As 14-3-3 avidity towards dimers increases and single bonds weaken, the inflection point at which increasing 14-3-3 concentration reduces activation moves right to higher [14-3-3]. **(b)** This effect is not due to the avidity change; modifying *K*_*double*_ alone does not move the inflection point. **(c)** Weakening the initial binding to any one site is sufficient to move the inflection point

## 4 Analytic Results

### 4.1 Dimer Derivative

As there exists an explicit solution to the model, it is possible to analytically find the values and derivatives of concentrations of various species. To understand the numerical results above, we found total dimer concentration,

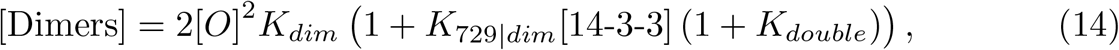

and took its derivative with respect to 14-3-3 concentration. The derivative is positive when two conditions hold. The first is that

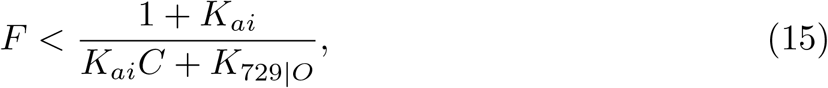

using the definitions

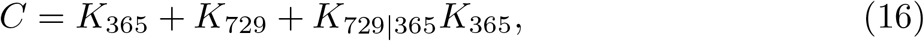

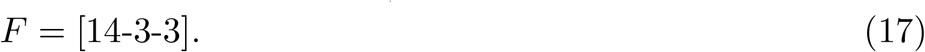

This requirement can be fulfilled as long as the system is not too heavily weighted towards monomers (by requiring sufficiently low *K*_729|*O*_ and *CK*_*ai*_), and can also be expressed as

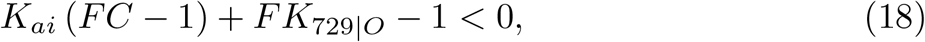

which is necessary for the next derivation.

The second condition is that

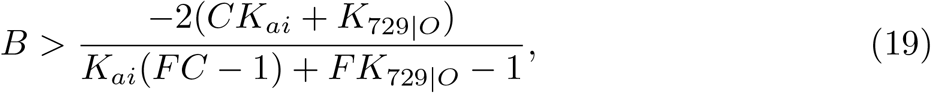

using the definition

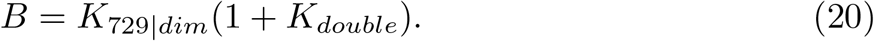

Using eqaution 18, this is equivalent to

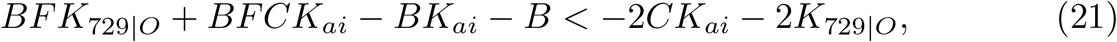

which reduces via

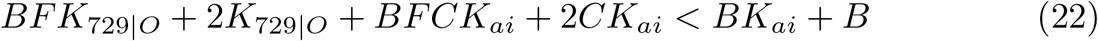

To

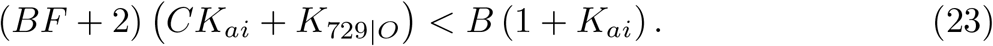

As total 14-3-3 increases monotonically with free 14-3-3, these conditions also describe when the derivative of dimer concentration with respect to total 14-3-3 is positive.

It is clear from Equation 23 that increasing 14-3-3 affinity for dimers (which increases *B*) is not sufficient to make 14-3-3 increase dimer concentration. To ensure the inequality holds, there must be either low free 14-3-3 (*F*) or low affinity for single bonds (*C* and *K*_729|*O*_, as shown in Figure 4c). As the total cellular concentration of 14-3-3 is much higher than that of RAF (Kulak et al, 2014), this suggests that single binding is relatively weak. In addition, Equation 23 requires that monomer avidity (*K*_729|365_, as part of *C*) is not too high, or that it is balanced by low single affinities for monomers (*K*_365_, *K*_729_).

### 4.2 Constant Single Binding

A useful simplification to understand these results is to hold all single binding constants (*K*_365_, *K*_729_, *K*_729|*O*_, *K*_729|*dim*_) equal, with their value set to *K*_*single*_. If you expand Equation 15 using

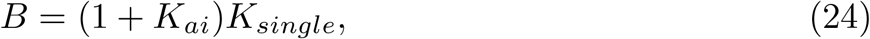

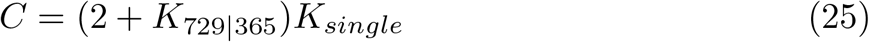

it gives

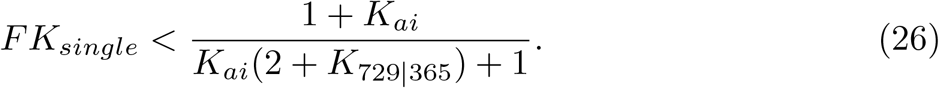

If this is stated in *K*_*D*_ terms (*K*_*D,single*_ = 1/*K*_*single*_), we find

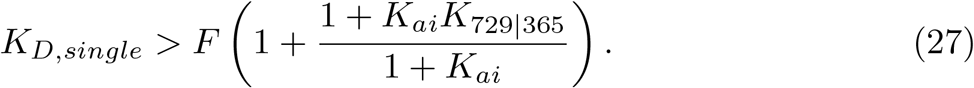

A similar substitution in Equation 23 gives the somewhat unwieldy

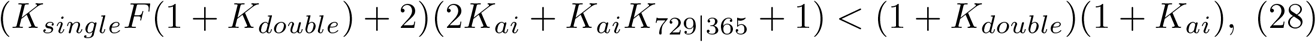

which is linear in *FK*_*single*_, so gives another condition where *K*_*D,single*_ is a rational expression multiplied by [14-3-3]. To ensure the result for *FK*_*single*_ is positive after the subtraction of the +2 term, a second condition is required:

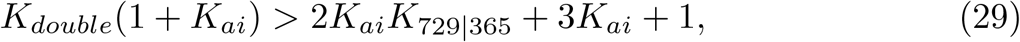

which serves as a lower bound for *K*_*double*_. Overall, we find that *K*_*D,single*_ has a lower bound of the greater of two rational expressions multiplied by [14-3-3], and *K*_*double*_ must at least

For a somewhat more concrete example of these bounds, consider the case of *K*_*ai*_ = Equation 23 then implies

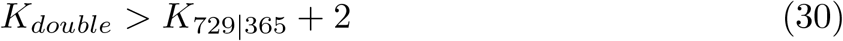

via Equation 29 and

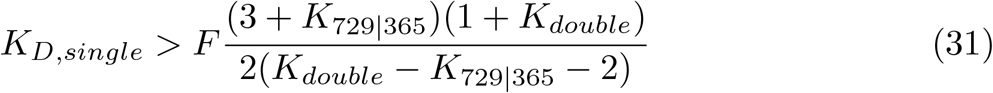

via Equation 28, while Equation 27 reduces to

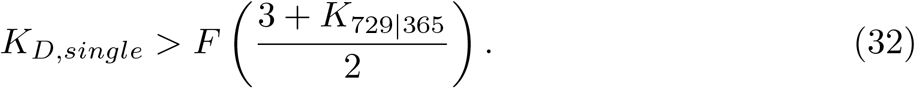

If we additionally set *K*_729|365_ = 1, this becomes

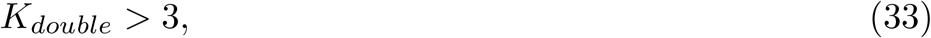

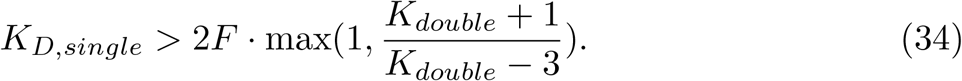

These results suggest *K*_*D,single*_ is significantly higher than [14-3-3], making single binding fairly weak regardless of the available 14-3-3 concentration.

### 4.3 Double-Binding Limit

To fulfill the conditions 23 and 18, which are required for the observed behavior that RAF dimerization increases as a result of overexpression of 14-3-3, it is necessary for 14-3-3–RAF binding at any single phosphosite to relatively weak, but significant avidity at the second binding site is still permitted. Therefore, it is informative to take the limit in which only double binding can occur. At this limit,

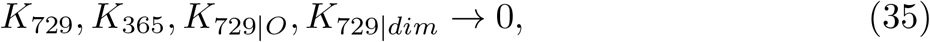

while the products *K*_729|*dim*_ · *K*_*double*_ and *K*_365_ · *K*_729|365_ are held constant. For this derivation, it is more convenient to use *K*_*D*_ notation than *K*_*A*_ notation (where *K*_*D*_ = 1/*K*_*A*_), so let

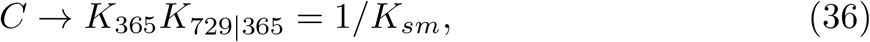

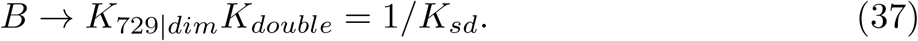

From there, condition 23 reduces to

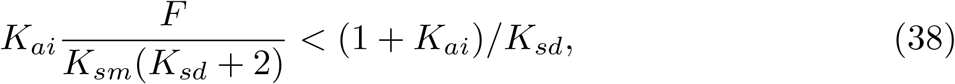

letting us further show

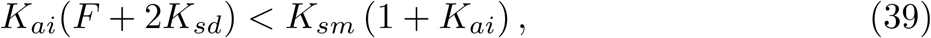

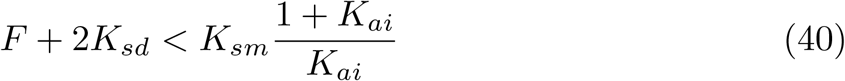

identically to the analogous condition in Mendiratta et al (2021) (equations unpublished). That model in the absence of drugs is the limit of this model when only double binding is significant, as is required for a positive derivative of dimer concentration with respect to 14-3-3.

## 5 Discussion

Given the experimental result that increased concentration of 14-3-3 increases RAF dimerization (Mendiratta et al, 2021), this model suggests that 14-3-3 binding to each individual phosphoserine in RAF is likely to be relatively weak. This may help clear up the highly variable results of previous simulations and experimental measurements of the affinity (Ghosh et al, 2015; Tzivion et al, 1998; Zhang et al, 2021). It also validates the simplifying approximation that 14-3-3 only binds in cases where both phosphosites are available, which we used in previous RAF modeling (Mendiratta et al, 2021) and may reuse in future work.

This system resembles Ercolani and Schiaffino’s model of chelate cooperativity (Ercolani and Schiaffino, 2011), (similar to avidity) but considers different limiting cases. Like our work, they find situations where one binding state will tend to become favored at sufficiently high concentrations due to 1:1 vs 2:1 ratios, specifically that at high enough concentrations of BB the “cyclic” double-binding (comparable to monomers) should be disfavored. (Their AA receptor is analogous to 14-3-3 and their BB ligand is analogous to RAF.) However they don’t consider the high AA receptor case, and since their results are in terms of free ligand concentration they don’t include the effects of receptor concentration on the free ligand. This makes sense when considering a ligand-receptor system, as they did, but does not necessarily generalize to other protein-protein interactions. In our model, 14-3-3 has a substantially higher concentration than RAF, and our work was partly motivated by experiments involving 14-3-3 overexpression. The effect of 14-3-3 concentration on free RAF is necessary for the stoichiometric preference for monomers at high 14-3-3 concentrations which we observe here, and our work goes on to show the conditions under which this stoichiometric pressure becomes dominant.

Many questions remain on RAF–14-3-3 interactions. While this result qualitatively constrains the strength of double-vs single-binding of 14-3-3 to RAF, due to the number of free parameters without reliable experimental measurement we cannot assign any specific value to each affinity without assumptions about other values. In addition, as this is not a complete model of RAF activation, any such value derived from this model would not be a physical equilibrium constants between two strictly defined states, but rather a composite over some collection of states. For instance, the value of *K*_*ai*_ in this model implicitly includes the effects of RAS binding (as might the 14-3-3–closed monomer affinities, depending the details of the relationship between RAS–RAF binding and RAF–14-3-3 binding (Martinez Fiesco et al, 2022)).

There is also considerable work to be done in RAF modeling more generally. We know of a wide variety of processes involved in RAF activation, including dimerization (and associated internal phosphorylation events), phosphorylation and dephosphorylation of S365 (the latter via the recently-imaged PP1C-SHOC2-MRAS complex (Bonsor et al, 2022; Hauseman et al, 2022; Kwon et al, 2022; Liau et al, 2022)), conformational changes, and RAS binding. The development of larger scale models that include additional steps of RAF regulation and additional binding partners is limited in part by uncertainties in mechanisms of RAF regulation. As models become more complex, numerical results become increasingly important, and the fact that many parameters have not been measured adds additional complexity to developing such models. We believe that focused studies like this that can facilitate the development of larger scope models both by justifying simplifications and by constraining which portions of parameter space better correspond with empirical observations.

## Acknowledgments

The authors thank Gaurav Mendiratta, Jodie Jepson, Craig Allen, Melinda Tong, Michael Jones, and the members of the Stites laboratory for helpful discussions and/or for comments on the manuscript.

## Declarations

This work was supported in part by NIH DP2AT011327.

## Appendix A Definitions

All concentrations are in molarity, all time units are seconds.

### A.1 Concentrations

Definitions of the concentration variables used in the model:

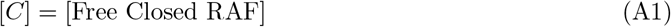

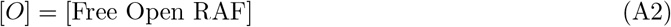

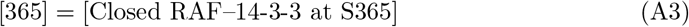

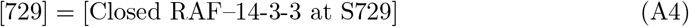

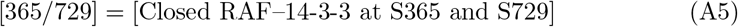

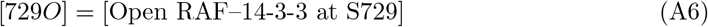

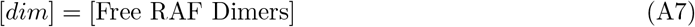

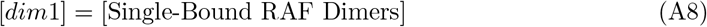

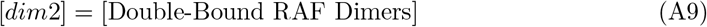

### A.2 Equilibrium Constants

Definitions of the equilibrium constants of each reaction in the model:

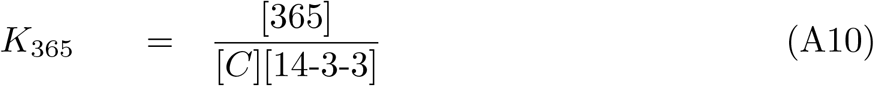

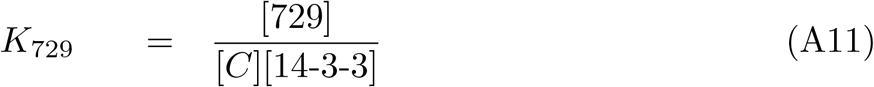

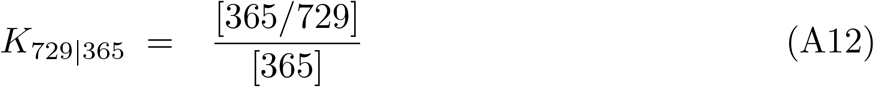

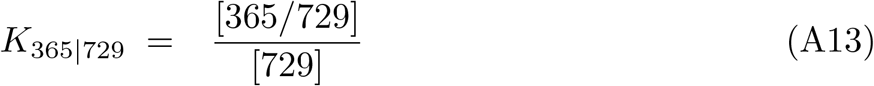

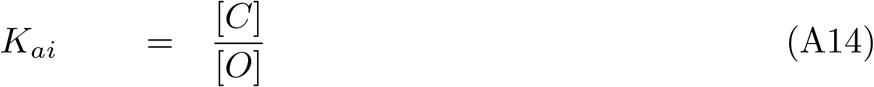

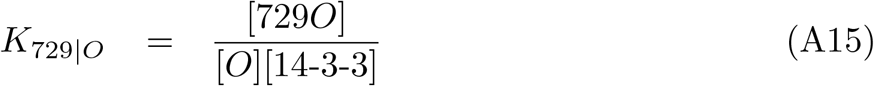

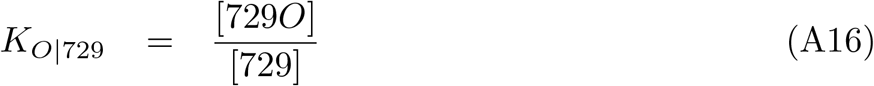

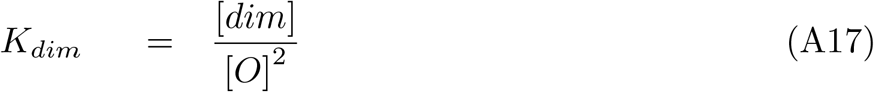

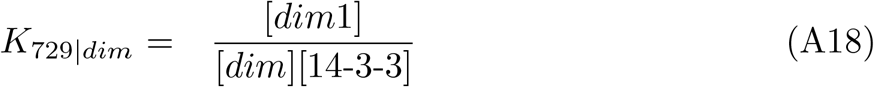

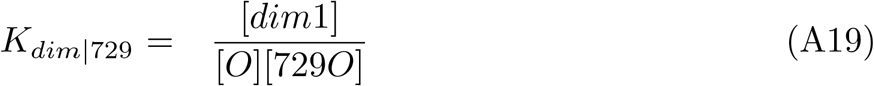

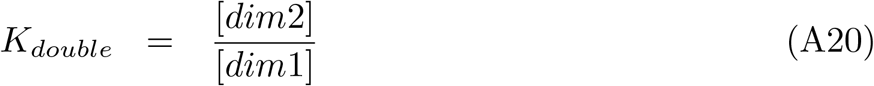

Note: — can be read as “given”.

### A.3 Detailed Balance

The detailed balance relations listed below are required to ensure that each state of the system has a consistent internal energy, and energy is not gained or lost as a molecule moves along a cycle to return to its initial state.

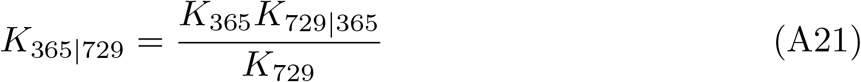

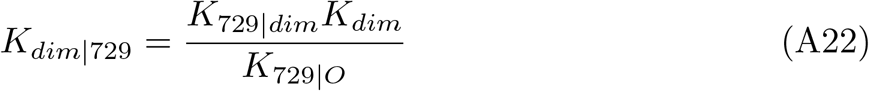

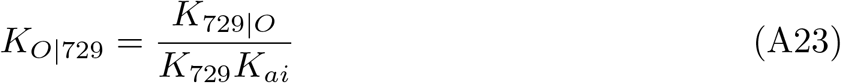

## Appendix B Default Values

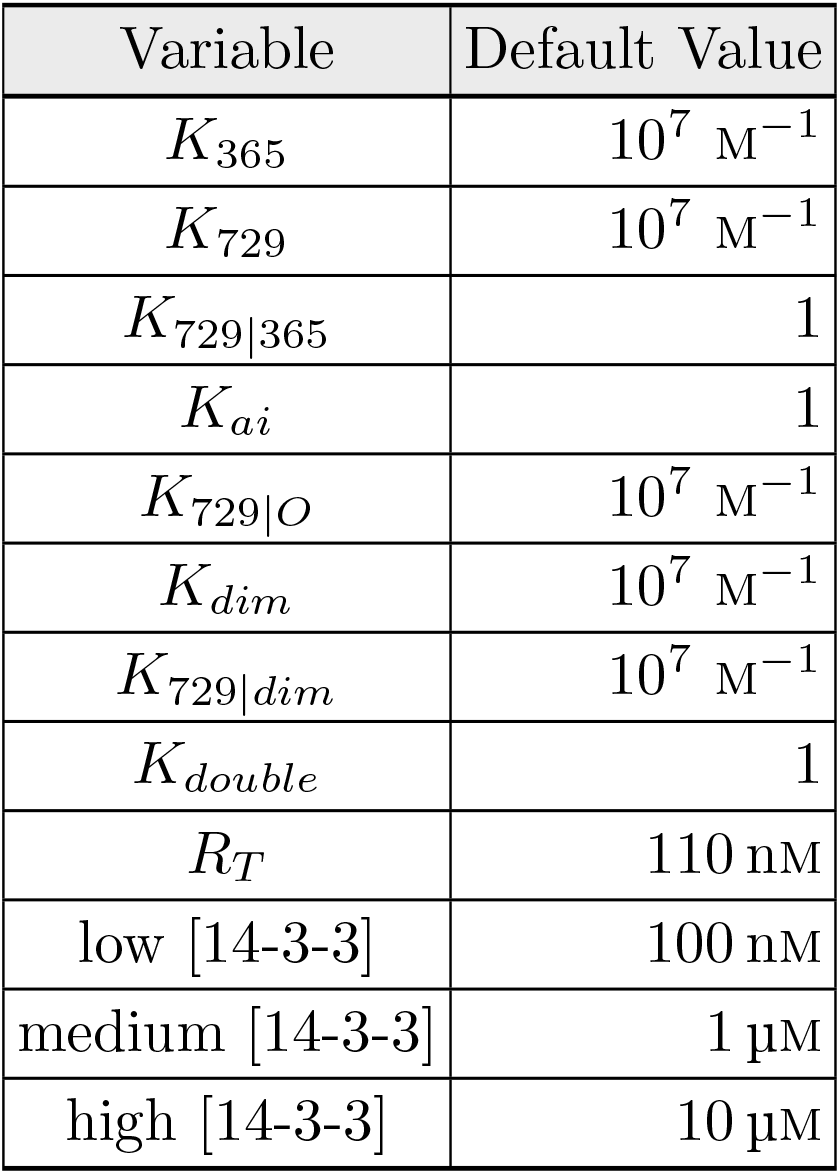

All binding constants above are order-of-magnitude estimates of plausible values for protein-protein binding, rather than measured values. The concentration of RAF was derived from the number of counts found in Kulak et al (2014) for HeLa cells, counting all isoforms of RAF. They were scaled to the necessary cell volume to give a total RAS concentration matching Fujioka et al (2006), with the assumption that KRAS (which Kulak did not measure) makes up half of total RAS. 14-3-3 concentrations were determined using the same process, under the assumption that ∼1/3, 1/30, and 1/300 of 14-3-3 ζ and *ϵ* were available for binding in the high, medium, and low cases respectively. Those isoforms were chosen based on the finding of Martinez Fiesco et al (2022) that 75% of 14-3-3 bound to RAF was of those isoforms.

## References

Bonsor DA, Alexander P, Snead K, et al (2022) Structure of the SHOC2–MRAS– PP1C complex provides insights into RAF activation and Noonan syndrome. Nature Structural & Molecular Biology 29(10):966–977. 10.1038/s41594-022-00841-4

Ercolani G, Schiaffino L (2011) Allosteric, Chelate, and Interannular Cooperativity: A Mise au Point. Angewandte Chemie International Edition 50(8):1762–1768. 10.1002/anie.201004201

Erlendsson S, Teilum K (2021) Binding Revisited—Avidity in Cellular Function and Signaling. Frontiers in Molecular Biosciences 7. 10.3389/fmolb.2020.615565

Fröhlich F, Gerosa L, Muhlich J, et al (2023) Mechanistic model of MAPK signaling reveals how allostery and rewiring contribute to drug resistance. Molecular Systems Biology 19(2):e10988. 10.15252/msb.202210988

Fu H, Subramanian RR, Masters SC (2000) 14-3-3 Proteins: Structure, Function, and Regulation. Annual Review of Pharmacology and Toxicology 40(1):617–647. 10.1146/annurev.pharmtox.40.1.617

Fujioka A, Terai K, Itoh RE, et al (2006) Dynamics of the Ras/ERK MAPK Cascade as Monitored by Fluorescent Probes*. Journal of Biological Chemistry 281(13):8917–8926. 10.1074/jbc.M509344200

Ghosh A, Ratha BN, Gayen N, et al (2015) Biophysical Characterization of Essential Phosphorylation at the Flexible C-Terminal Region of C-Raf with 14-3-3? Protein. PLOS ONE 10(8):e0135976. 10.1371/journal.pone.0135976

Hauseman ZJ, Fodor M, Dhembi A, et al (2022) Structure of the MRAS–SHOC2– PP1C phosphatase complex. Nature 609(7926):416–423. 10.1038/s41586-022-05086-1

Hu J, Stites EC, Yu H, et al (2013) Allosteric Activation of Functionally Asymmetric RAF Kinase Dimers. Cell 154(5):1036–1046. 10.1016/j.cell.2013.07.046

Kholodenko BN (2015) Drug Resistance Resulting from Kinase Dimerization Is Rationalized by Thermodynamic Factors Describing Allosteric Inhibitor Effects. Cell Reports 12(11):1939–1949. 10.1016/j.celrep.2015.08.014

Kulak NA, Pichler G, Paron I, et al (2014) Minimal, encapsulated proteomic-sample processing applied to copy-number estimation in eukaryotic cells. Nature Methods 11(3):319–324. 10.1038/nmeth.2834

Kwon JJ, Hajian B, Bian Y, et al (2022) Structure–function analysis of the SHOC2– MRAS–PP1C holophosphatase complex. Nature 609(7926):408–415. 10.1038/s41586-022-04928-2

Lavoie H, Therrien M (2015) Regulation of RAF protein kinases in ERK signalling. Nature Reviews Molecular Cell Biology 16(5):281–298. 10.1038/nrm3979

Liau NPD, Johnson MC, Izadi S, et al (2022) Structural basis for SHOC2 modulation of RAS signalling. Nature 609(7926):400–407. 10.1038/s41586-022-04838-3

Martinez Fiesco JA, Durrant DE, Morrison DK, et al (2022) Structural insights into the BRAF monomer-to-dimer transition mediated by RAS binding. Nature Communications 13(1):486. 10.1038/s41467-022-28084-3

Mendiratta G, Stites E (2023) Theoretical analysis reveals a role for RAF conformational autoinhibition in paradoxical activation. eLife 12:e82739. 10.7554/eLife.82739

Mendiratta G, Abbott K, Li YC, et al (2021) RAF conformational autoinhibition and 14-3-3 proteins promote paradoxical activation. 10.1101/849489

Morrison DK, Heidecker G, Rapp UR, et al (1993) Identification of the major phosphorylation sites of the Raf-1 kinase. Journal of Biological Chemistry 268(23):17309–17316. 10.1016/S0021-9258(19)85336-X

Muslin AJ, Tanner JW, Allen PM, et al (1996) Interaction of 14-3-3 with Signaling Proteins Is Mediated by the Recognition of Phosphoserine. Cell 84(6):889–897. 10.1016/S0092-8674(00)81067-3

Park E, Rawson S, Li K, et al (2019) Architecture of autoinhibited and active BRAF–MEK1–14-3-3 complexes. Nature 575(7783):545–550. 10.1038/s41586-019-1660-y

Rukhlenko OS, Khorsand F, Krstic A, et al (2018) Dissecting RAF Inhibitor Resistance by Structure-based Modeling Reveals Ways to Overcome Oncogenic RAS Signaling. Cell Systems 7(2):161–179.e14. 10.1016/j.cels.2018.06.002

Simanshu DK, Morrison DK (2022) A Structure is Worth a Thousand Words: New Insights for RAS and RAF Regulation. Cancer Discovery 12(4):899–912. 10.1158/2159-8290.CD-21-1494

Tzivion G, Luo Z, Avruch J (1998) A dimeric 14-3-3 protein is an essential cofactor for Raf kinase activity. Nature 394(6688):88–92. 10.1038/27938

Zhang M, Jang H, Li Z, et al (2021) B-Raf autoinhibition in the presence and absence of 14-3-3. Structure 29(7):768–777.e2. 10.1016/j.str.2021.02.005

